# OptSurvCutR: Validated Cut-point Selection for Survival Analysis

**DOI:** 10.1101/2025.10.08.681246

**Authors:** Payton TO Yau

## Abstract

The stratification of subjects based on continuous predictors is a common yet challenging task in time-to-event analysis, particularly when relationships are non-linear and require multiple thresholds. Arbitrary cut-point selection inflates Type I error rates and produces biased effect estimates. This manuscript presents OptSurvCutR (Optimal Survival Cut-Points in R), addressing this challenge through three key logical processes: (1) data-driven determination of the optimal number of cut-points using information criteria; (2) simultaneous identification of multiple thresholds via a genetic algorithm; and (3) integrated bootstrap validation to assess cut-point stability and control false discovery risk. First, the *find_cutpoint_number()* function determines the optimal model complexity by comparing metrics such as AIC, AICc, or BIC, while optionally adjusting for covariates. Second, *find_cutpoint()* identifies the precise threshold locations by optimising survival metrics (e.g., log-rank statistic, hazard ratio) using either an exhaustive systematic search or an efficient genetic algorithm. Finally, *validate_cutpoint()* assesses the robustness of identified thresholds by generating 95% confidence intervals through bootstrap resampling. We demonstrate the package’s complete workflow using two case studies: a plant science example modelling the non-monotonic effect of temperature on rapeseed germination, and a clinical bioinformatics analysis stratifying colorectal cancer patients by a microbial biomarker after adjusting for clinical covariates. These examples illustrate how OptSurvCutR uncovers complex survival patterns often missed by traditional dichotomisation. The package provides a transparent, extensible framework for validated cut-point selection, making it broadly applicable for researchers working with time-to-event data. Source code and documentation are freely available at https://github.com/paytonyau/OptSurvCutR.

## BACKGROUND

The identification of meaningful thresholds within continuous data is a ubiquitous task in data-driven research, from clinical diagnostics to environmental and food science. The conversion of a continuous variable into a categorical one by defining an optimal cut-point is a powerful method for risk stratification and decision-making (Mayeux, 2004). In high-throughput omics data, such as gene expression, for example, it often relies on this cut-off approach to stratify patients into low- and high-risk groups for survival outcomes.

However, identifying a truly optimal and robust cut-point is a non-trivial statistical challenge. The choice of threshold is so influential that different cut-points can lead to contradictory conclusions from the same dataset, creating a high risk of “data-dredging bias” if the method is not transparently reported and justified (Tustumi, 2022). Foundational simulation studies have long confirmed that outcome-oriented cut-point selection, if not properly validated, can produce biased estimates of effect size and substantially inflated Type I error rates (Altman et al., 1994; Faraggi & Simon, 1996). Despite these well-documented dangers, the need for accessible and statistically sound tools persists. For survival data, the “maximally selected rank statistic”—which identifies the cut-point that maximises the separation between survival curves—is a well-established outcome-oriented approach (Mandrekar et al., 2003; Meyers & Mandrekar, 2015).

A further complication is that biological and environmental systems rarely adhere to simple linear relationships. The limitations of standard parametric models in this context have been highlighted by health technology assessment bodies, which recommend the use of more flexible methods to adequately capture complex hazard functions, particularly for long-term extrapolation (Rutherford et al., 2020). A single cut-point is often insufficient for capturing non-monotonic, J-shaped, S-shaped, and U-shaped relationships, such as when both very low and very high levels of a biomarker (i.e., temperature and pH) are associated with increased risk (Royston et al., 2006). In these cases, forcing such complex relationships into a simple high/low dichotomy can lead to significant patient misclassification and obscure the true underlying biological patterns. Identifying multiple thresholds to define low-, intermediate-, and high-risk groups is therefore essential for accurately modelling such complex systems. Furthermore, any identified thresholds must be evaluated for their prognostic value independent of known clinical or demographic confounders, such as patient age or disease stage. Failure to adjust for covariates can lead to spurious associations where a biomarker appears significant only because it correlates with another established risk factor.

While several R packages exist to address this challenge, they are often specialised for specific scenarios. Packages like *survminer* provide excellent tools for finding a single optimal cut-point in survival contexts (Alboukadel Kassambara et al., 2024). Foundational methods for outcome-based biomarker assessment were established by early graphical bioinformatics tools like X-tile, which was computationally limited to finding a maximum of two cut-points (Camp et al., 2004), and later by now-defunct web applications like Cutoff Finder (Budczies et al., 2012). More recent statistical approaches have also been described for determining both the optimal number and location of cut-points, for instance, by using the Akaike Information Criterion (AIC) to select the best model (Chang et al., 2017; Chen et al., 2019). Despite the power of these individual tools, a significant methodological gap remains for a single, well-integrated R package that provides a complete and flexible workflow—from determining the appropriate number of cuts to finding their locations and validating the result—within a reproducible software environment.

To address these limitations, we have developed *OptSurvCutR*, a versatile R package designed specifically for robust cut-point analysis in a time-to-event context. *OptSurvCutR* consolidates the entire analytical process into a single toolkit, offering (1) a data-driven function to determine the optimal number of cut-points, (2) flexible algorithms for identifying both single and multiple thresholds, (3) an integrated validation step to ensure the robustness of the findings, and (4) support for parallel processing to accelerate computationally intensive tasks.

A critical but often-overlooked practical concern is determining adequate sample size for reliable cut-point identification. While theoretical guidance exists for single cut-points, the sample size requirements for multiple thresholds remain poorly defined. This manuscript provides empirical observations on sample size adequacy through case studies and discusses practical limitations in the supplementary materials (need to integrate).

## IMPLEMENTATION

### Package Architecture and Workflow

The *OptSurvCutR* package is implemented in the R statistical programming language and is designed to provide a cohesive and logical workflow for cut-point analysis in a survival context. The workflow is structured to guide a researcher from initial data exploration to final model validation and is centred around three core functions, which are summarised in **Figure 1**: *find_cutpoint_number(), find_cutpoint()*, and *validate_cutpoint()*. These functions employ one of two distinct search algorithms for optimisation: a “systematic” search (Altman et al., 1994), and a “genetic” algorithm (Mebane Jr & Sekhon, 2011). The systematic search exhaustively evaluates all possible thresholds, guaranteeing a globally optimal solution but becoming computationally infeasible for multiple cut-points. In contrast, the genetic algorithm uses a heuristic evolutionary approach to efficiently search the vast parameter space, making it a practical and powerful tool for identifying multiple thresholds in larger datasets.

**Figure 1.**
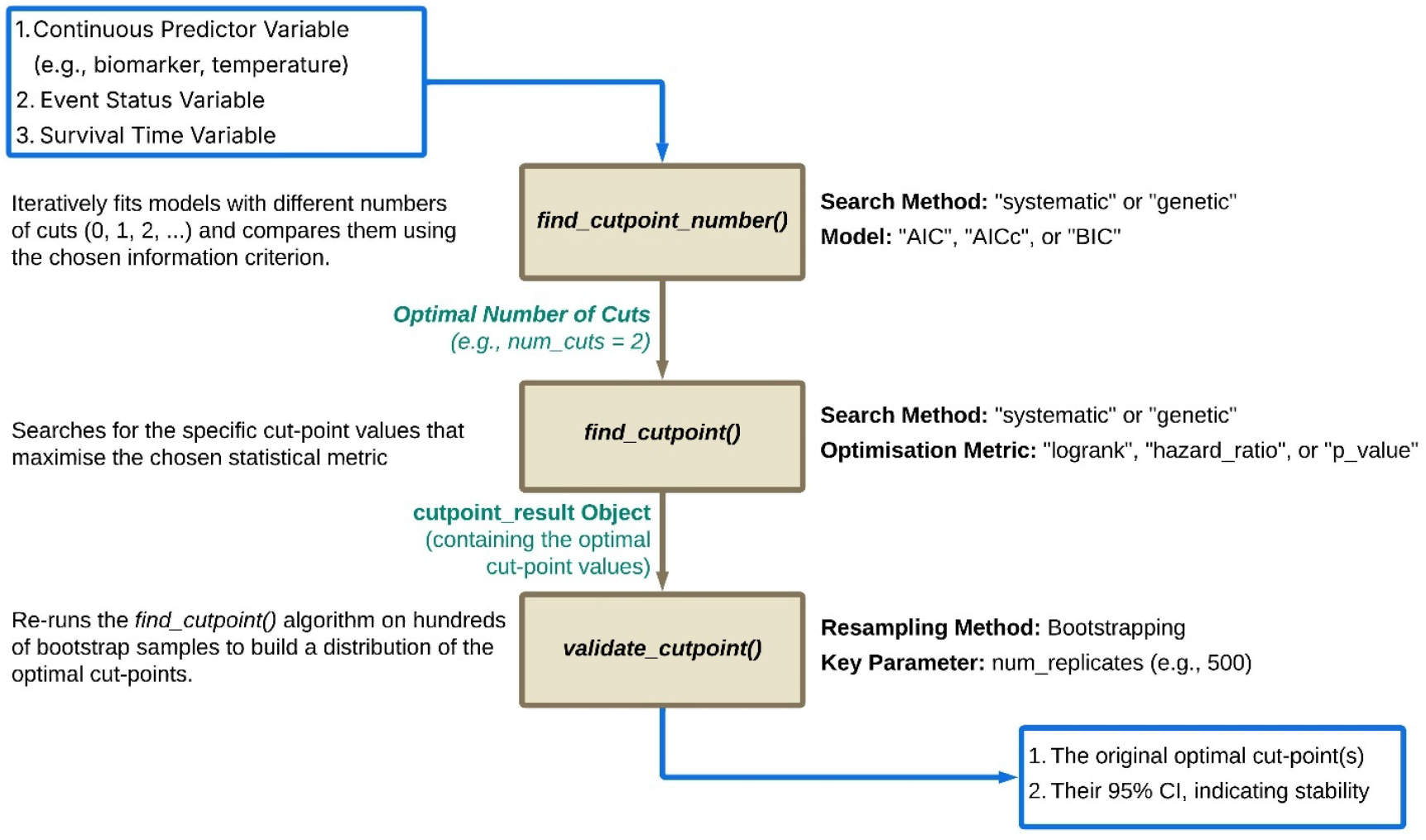
The OptSurvCutR analysis workflow. The flowchart outlines the three core functions and their recommended use in a typical analysis. AIC, Akaike Information Criterion; AICc, Corrected Akaike Information Criterion; BIC, Bayesian Information Criterion.

## CORE FUNCTIONS AND ALGORITHMS

### Step 1: Determining the Number of Cut-points with *find_cutpoint_number()*

This function serves as the initial, exploratory step to help researchers make a data-driven decision on the most plausible number of cut-points. It leverages the search algorithms to compare models with different numbers of thresholds (e.g., 0 vs. 1 vs. 2 cuts), optionally adjusting for specified covariates, and provides statistical evidence using information criteria. This allows for the selection of the optimal number of groups needed to explain the predictor’s effect *after* accounting for the influence of potential confounders. The function supports three information criteria (**Table 1**), each with distinct statistical properties and practical implications:

- **AIC**: Estimates out-of-sample prediction error via Kullback-Leibler divergence. Tends to select more complex models; recommended for smaller datasets (n < 100) (Akaike, 1974).
- **AICc**: Corrected AIC for small samples. Strongly recommended when n/(k+1) < 40, where k is the number of parameters (Hurvich & Tsai, 1989).
- **BIC**: Derives from Bayesian model averaging under uniform priors. Penalises complexity more heavily than AIC; recommended for larger datasets and when sparsity is desired (Gideon Schwarz, 1978).

**Table 1.** Information Criteria for Model Selection.

| Criterion | Formula | Components |
| --- | --- | --- |
| **Akaike Information Criterion (AIC)** | $-2 \cdot \log(L) + 2k$ | $L$ = Maximised likelihood<br>$k$ = Number of parameters |
| **Corrected Akaike Information Criterion (AICc)** | $AIC + \frac{2k(k+1)}{n-k-1}$ | $n$ = Sample size<br>$L$ and $k$ as above |
| **Bayesian Information Criterion (BIC)** | $-2 \cdot \log(L) + k \cdot \log(n)$ | $L$, $k$, and $n$ as above |

For most survival analyses, we recommend BIC as the default, with AICc as a secondary option for n < 100. The function computes Akaike weights (exp(−ΔIC/2)) to quantify relative model plausibility (Supplementary Information Section S2).

### Step 2: Identifying Cut-point Locations with *find_cutpoint()*

This is the main workhorse function for identifying the precise location of the optimal cut-point(s). It uses one of two search algorithms—either a systematic or genetic search—to find the cut-point(s) that optimise a user-selected survival metric, while allowing for adjustment via the covariates argument. This crucial feature enables researchers to assess whether the prognostic value of the cut-points derived from the continuous predictor is independent of other known risk factors (e.g., age, sex, tumour stage). The available optimisation criteria are based on established statistical methods:

- **“logrank”**: This criterion finds the cut-point that maximises the chi-squared statistic from the standard non-parametric log-rank test (Mantel, 1966). The test compares the observed (O) and expected (E) number of events across groups at each event time.
- **“hazard_ratio”**: This criterion finds the cut-point that maximises the Hazard Ratio (HR) derived from a Cox Proportional-Hazards model (Cox, 1972). The HR quantifies the magnitude of the difference in risk between the groups created by the cut-point, with a higher value indicating a more clinically significant separation.
- **“p_value”**: This criterion seeks the cut-point that minimises the p-value from the log-rank test component of a Cox Proportional-Hazards model (Cox, 1972). This method directly optimises for the highest level of statistical significance in the difference between the survival curves of the groups.

### Step 3: Assessing Stability with *validate_cutpoint()*

After a cut-point has been identified, this function assesses its stability and robustness. It performs a non-parametric bootstrap resampling procedure (Efron, 1979). This process involves creating a large number of new datasets by sampling subjects with replacement from the original data. To ensure a fair validation, the entire *find_cutpoint()* algorithm, using the original settings (e.g., method and criterion), is then re-applied to each of these bootstrap samples. The resulting collection of newly discovered optimal cut-points is used to build an empirical distribution, which provides a direct assessment of how sensitive the optimal cut-point is to sampling variability. This distribution is then used to calculate 95% confidence intervals, which indicate the reliability of the chosen threshold.

The package also includes a suite of S3 methods for *print(), summary()*, and *plot()* to ensure that the results from each step are user-friendly and easy to interpret visually.

### Software Implementation and Availability

The *OptSurvCutR* package is implemented in R (R Core Team, 2025). Its core statistical functionality relies on the *survival* package for time-to-event modelling (Therneau, 2024) and the *rgenoud* package for the genetic algorithm optimisation (Mebane Jr & Sekhon, 2011). For computational efficiency, parallel processing is implemented via the *foreach* and *doParallel* packages for the bootstrap validation step, while the search algorithms utilise sequential processing to ensure numerical stability (Microsoft Corporation & Weston, 2022). Data manipulation and visualisation are handled by packages from the *tidyverse* ecosystem (Wickham et al., 2019), primarily *dplyr* (Wickham et al., 2014) and *ggplot2* (Wickham, 2016), with survival-specific plots enhanced by *survminer* (Kassambara et al., 2016). User-facing messages and progress bars are generated by the *cli* package (Csárdi, 2017). Reproducibility for the stochastic genetic algorithm is ensured through base R’s random seed generation. The package, including all source code and documentation, is freely available from GitHub at https://github.com/paytonyau/OptSurvCutR.

## CASE STUDIES

### Case Study 1: Identifying Temperature Thresholds for Rapeseed Germination

To demonstrate the full exploratory workflow of *OptSurvCutR* in a time-to-event context, we analysed a simulated dataset inspired by research on the germination of rapeseed (*Brassica napus L*.), where germination time is expected to have a non-monotonic relationship with temperature (Haj Sghaier et al., 2022). To create a realistic dataset for analysis, we simulated data in two parts to mimic the findings of the original study. First, we generated data based on the seven constant temperature points reported in the manuscript. Then, to create a more continuous predictor variable for finding optimal thresholds, we interpolated the germination parameters for six intermediate temperature points and generated a second part of the dataset. The combination of these simulated datasets is used to demonstrate the capabilities of the package (**Supplementary information 1**).

First, an exploratory analysis was performed using *find_cutpoint_number()* to determine the most plausible number of cut-points. A comparison of models with 0 to 4 cuts showed that the BIC was minimised for a model with 3 cut-points, providing statistical support for stratifying the data into four distinct temperature zones (**Figure 2A**).

**Figure 2.**
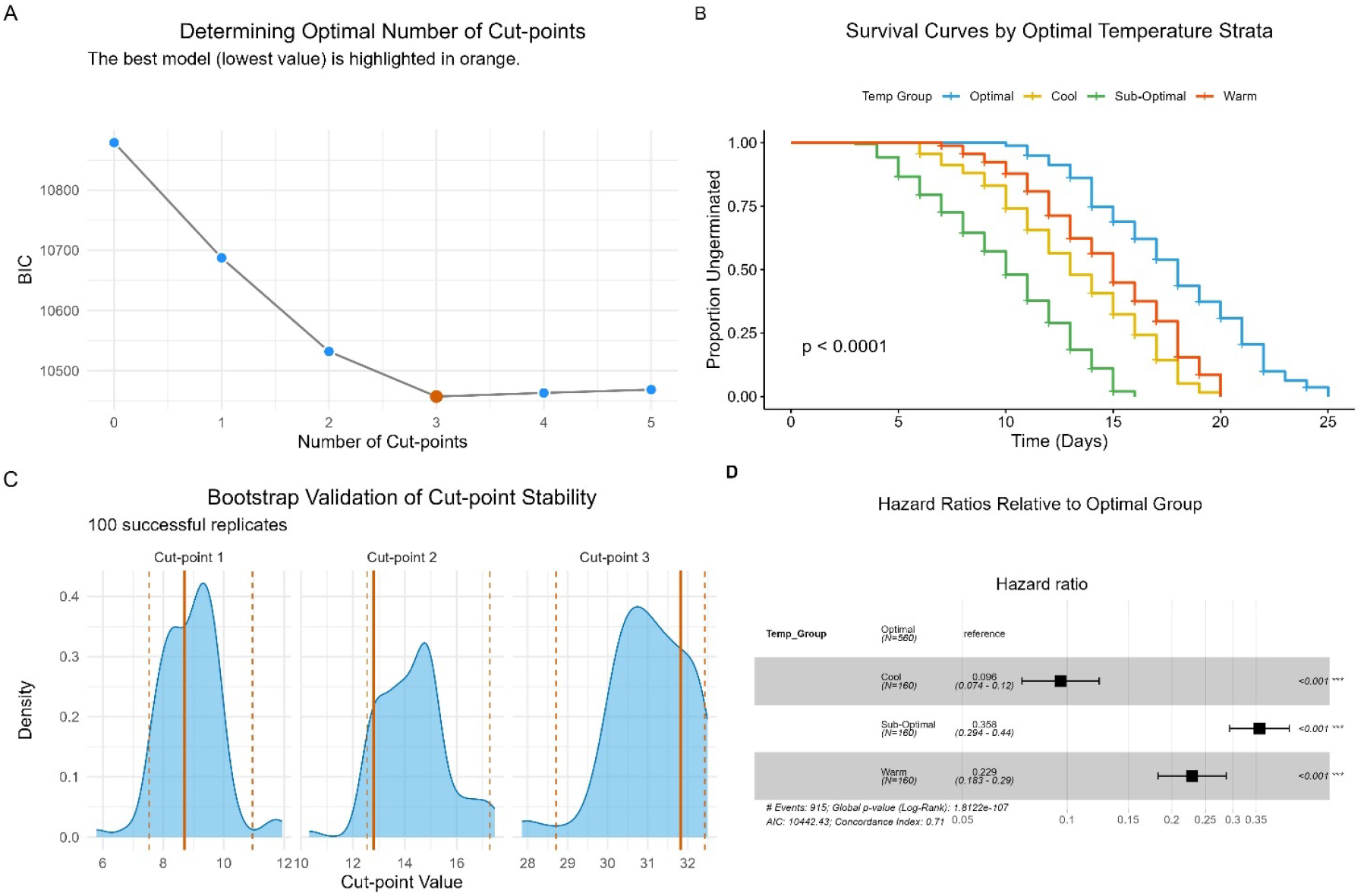
A comprehensive workflow demonstrating the statistical identification, validation, and interpretation of optimal temperature thresholds for rapeseed germination time. (A) Optimal Model Selection. Bayesian Information Criterion (BIC) scores plotted against the number of proposed cut-points. The model with 3 cut-points was selected as optimal due to the lowest BIC score, justifying the stratification of the data into four distinct groups. **(B) Survival Analysis:** Kaplan-Meier curves illustrating the time-to-germination profiles for the four statistically derived temperature groups: “Cool”, “Sub-Optimal”, “Optimal”, and “Warm”. The “Optimal” group exhibits the fastest germination and is set as the reference for comparison in the hazard ratio analysis. Shaded areas represent 95% confidence intervals, and the displayed p-value is from a log-rank test assessing the overall difference between the groups. **(C) Cut-point Stability Assessment:** Density plots showing the empirical distribution of the three optimal cut-points derived from 500 bootstrap replicates. The narrow peaks indicate high stability of the originally identified thresholds (solid vertical lines). **(D) Effect Size Quantification:** A forest plot displaying the Hazard Ratios (HR) and 95% confidence intervals from a Cox proportional-hazards model. The analysis quantifies the germination rate of the non-optimal groups relative to the “Optimal” reference group. Error bars represent 95% confidence intervals.

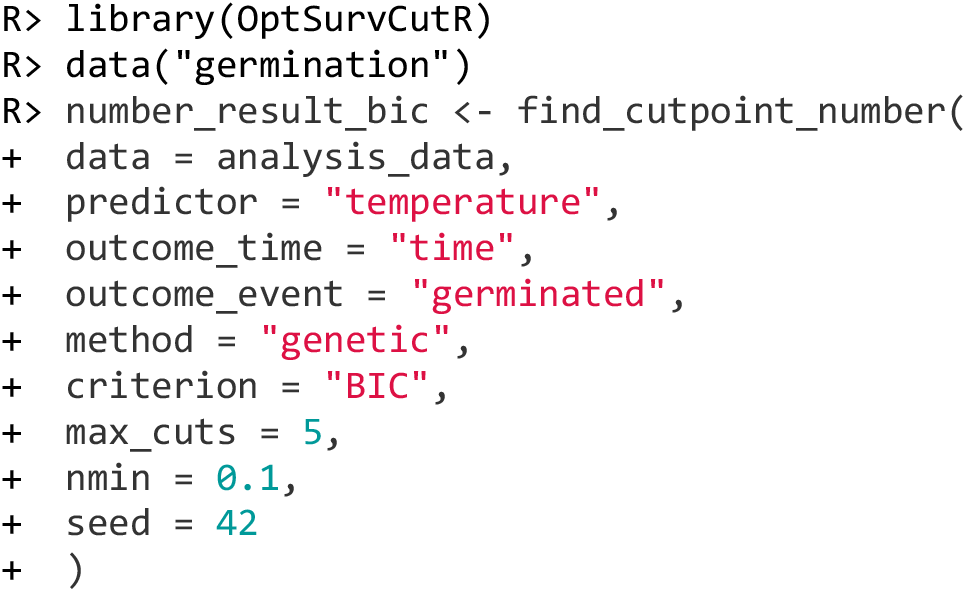

Next, *find_cutpoint()* was run with method = “genetic”, criterion = “logrank”, and num_cuts = 3 to identify the specific temperature thresholds. The algorithm identified three optimal cut-points at 8.7°C, 12.8°C, and 31.8°C. These thresholds stratified the seeds into four groups: “Cool” (≤8.7°C), “Sub-Optimal” (8.7°C-12.8°C), “Optimal” (12.8°C-31.8°C) and “Warm” (>31.8°C). The Kaplan-Meier analysis of the four groups showed that the “Optimal” group had the fastest germination rate, as indicated by an early and steep drop in its curve. In contrast, the “Cool”, “Sub-Optimal”, and “Warm” groups showed similar patterns of inhibited germination, with their curves dropping days after the “Optimal” group. It is important to note that this interpretation differs from a typical survival analysis; here, the “event” is successful germination, meaning a drop in the curve represents a positive outcome. The p-value (< 0.0001; **Figure 2B**) indicates the differences between these groups are highly statistically significant.

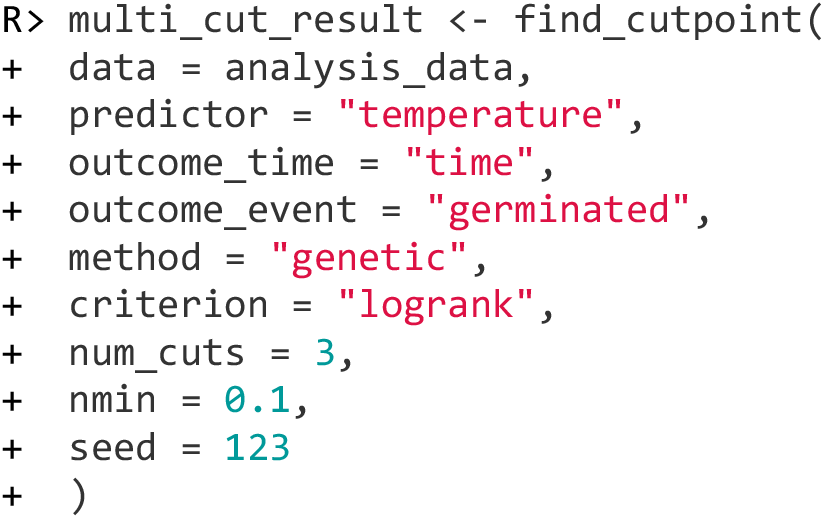

Finally, the stability of these three thresholds was assessed using *validate_cutpoint()* with 500 bootstrap replicates. The analysis yielded narrow 95% confidence intervals for all three cut-points, indicating that the identified thresholds were stable and not artefacts of the specific data sample (**Figure 2C**).

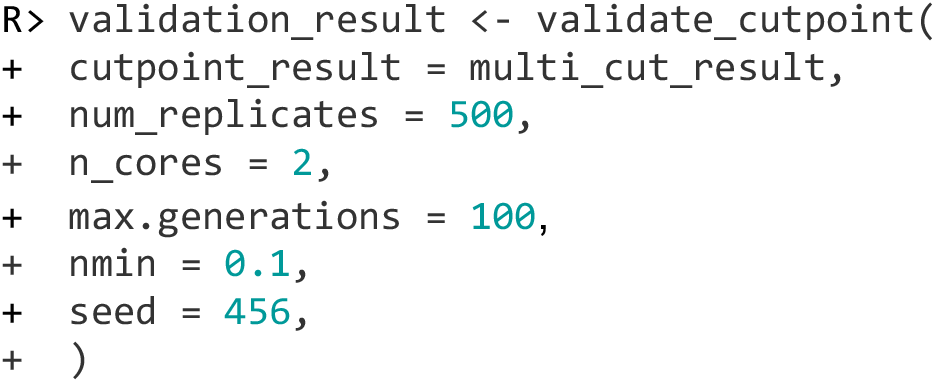

Analysis using a Cox proportional-hazards model showed that seeds in the “Cool”, “Sub-Optimal”, and “Warm” temperature groups had a significantly slower germination time compared to the “Optimal” reference group. The result was highly significant (*p* < 0.0001), as visualised in the corresponding forest plot (**Figure 2D**).

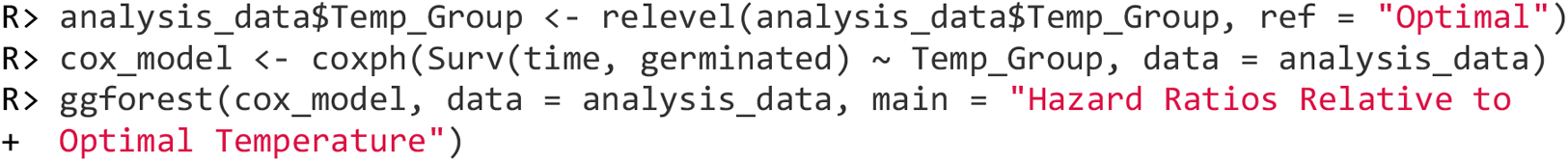

### Case Study 2: Stratifying Colorectal Cancer Patients by a Microbial Biomarker

To demonstrate the application of *OptSurvCutR* in a clinical bioinformatics context, we utilised a TCGA gut virome dataset in colorectal cancer (CRC) (Smyth et al., 2024). We aimed to identify the relative abundance of the genus Enterovirus in patients across several distinct survival groups for a 5-year overall survival analysis, with patient follow-up time censored at 60 months.

First, an exploratory study using *find_cutpoint_number()* was performed to identify the optimal number of prognostic groups. A comparison of models with 0 to 4 cuts indicated that a model with two cut-points was optimal, as determined by the BIC (BIC = 1251.69) with using genetic and a minimum of 20% of the total number for the group to avoid low numbers in any one of the groups (**Figure 3A**).

**Figure 3.**
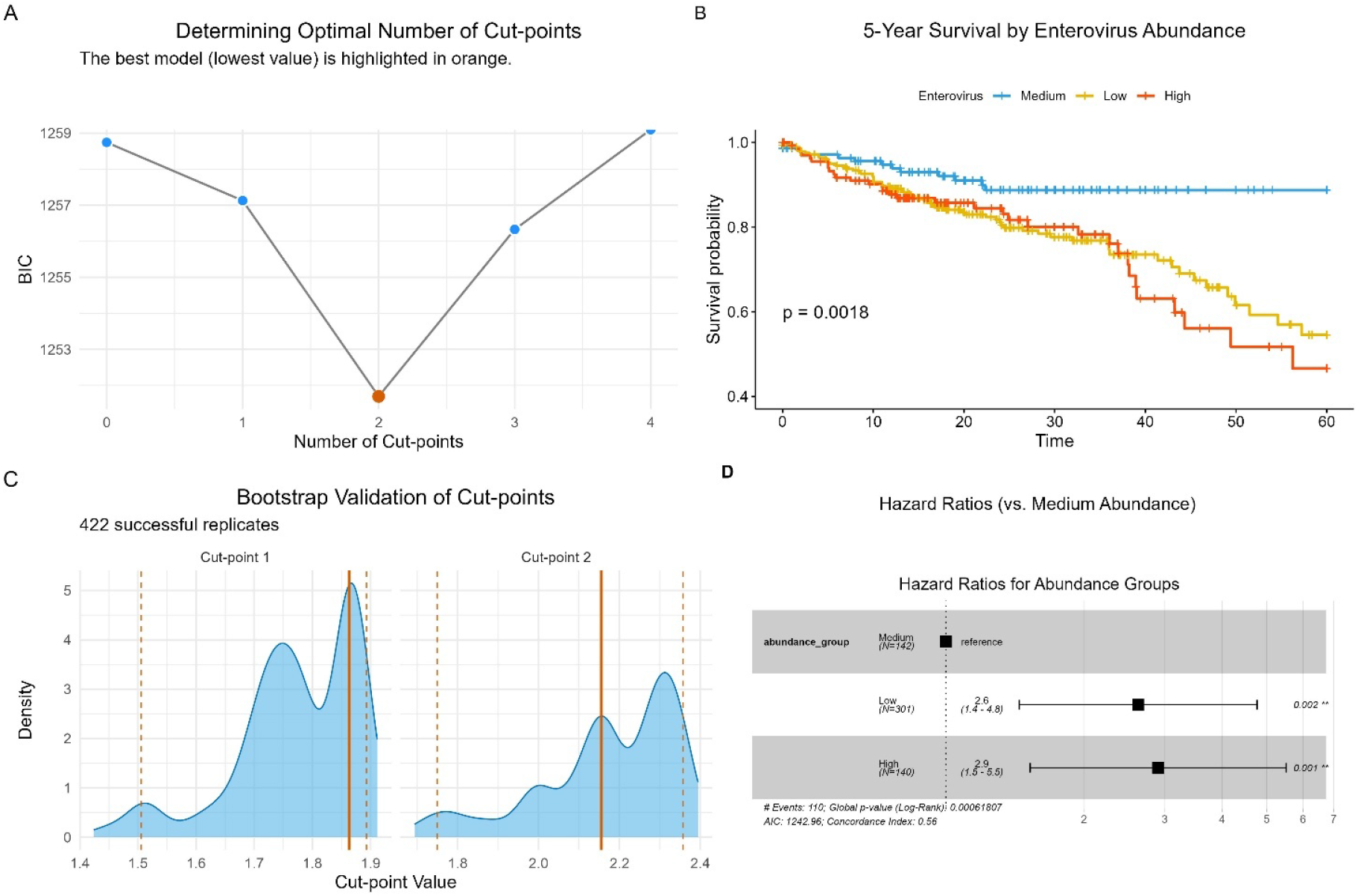
Complete workflow for the prognostic analysis of *Enterovirus* abundance in colorectal cancer. (A) Optimal Model Selection. Bayesian Information Criterion (BIC) analysis to determine the optimal number of prognostic groups based on *Enterovirus* abundance. The model with 2 cut-points (creating three groups) was selected due to the lowest BIC score. **(B) Survival Analysis:** Kaplan-Meier curves for 5-year overall survival for the three patient groups (“Low”, “Medium”, “High”). The analysis reveals a significant difference in survival profiles (log-rank p < 0.001), with the “Medium” abundance group showing the most favourable outcome. **(C) Cut-point Stability:** Density plots from a bootstrap analysis (500 replicates) assessing the stability of the two abundance thresholds. The narrow distributions confirm the robustness of the identified cut-points (solid vertical lines). **(D) Risk Quantification:** Forest plot displaying the Hazard Ratios (HR) for mortality from a Cox proportional-hazards model. The analysis quantifies the increased risk for the “Low” and “High” abundance groups relative to the “Medium” reference group. Error bars represent 95% confidence intervals.

Next, *find_cutpoint()* was executed with num_cuts = 2 to identify the precise abundance thresholds that best separated these three groups, optimising for the log-rank statistic. The two optimal cut-points were used to categorise patients into “Low”, “Medium”, and “High” Enterovirus genus abundance groups. A Kaplan-Meier analysis revealed a highly significant difference in 5-year overall survival among these three groups (log-rank, *p* = 0.0018; **Figure 3B**).

The stability of these thresholds was then assessed using *validate_cutpoint()* with 500 bootstrap replicates. The analysis yielded narrow 95% confidence intervals for both cut points (Cut 1: 1.506 – 1.893; Cut 2: 1.749 – 2.357) (**Figure 3C**). The narrow confidence intervals confirmed the stability of both thresholds, although their slight overlap suggests the boundary between the “Low”, “Medium”, and “High” groups may represent a continuous gradient of risk rather than an abrupt change.

A subsequent Cox proportional-hazards model was fitted to quantify the risk associated with each group, using the “Medium” abundance group as the reference. The “High” abundance group was associated with a significantly increased hazard of mortality (Hazard Ratio = 2.9; 95% CI, 1.5 - 5.5), while the “Low” group showed a risk profile similar to the reference group (Hazard Ratio = 2.6; 95% CI, 1.4 - 4.8)(**Figure 3D**).

These case studies demonstrated how *OptSurvCutR* can be used to uncover a complex, non-linear prognostic relationship, identifying a specific high-risk patient subgroup that would be obscured by traditional dichotomisation methods.

## Discussion

The identification of optimal cut-points is a common yet challenging task in data analysis, with significant implications for biomarker discovery and patient stratification. In this paper, we have introduced *OptSurvCutR*, a novel R package designed to provide a comprehensive and statistically robust workflow for this purpose in a time-to-event context. Through a case study using a simulated dataset inspired by real-world ecological research, we have demonstrated its key capabilities for identifying and validating multiple cut-points.

A primary advantage of *OptSurvCutR* is its integrated workflow, which guides the user from initial model selection to final validation. The validate_cutpoint() function is particularly critical, as it provides a direct, resampling-based solution to the well-documented problems of inflated significance and biased effect estimates that arise when optimal cut-points are selected without correction (Faraggi & Simon, 1996; Rota et al., 2015). This validation step is essential because a potential concern with any stochastic search method, like the genetic algorithm, is that a given result may be an artifact of a single ‘lucky’ random seed. Our package’s workflow is explicitly designed to address this. While the seed in *find_cutpoint()* ensures *reproducibility*, the *validate_cutpoint()* function provides the proof of *robustness*. By re-running the entire search algorithm on 500 bootstrap replicates, this function demonstrates that the identified thresholds are stable and not a spurious result of the initial search. The narrow 95% confidence intervals in both of our case studies (**Figure 2C** and **Figure 3C**) confirm that our findings are robust to sampling variability. The *find_cutpoint_number()* function addresses a critical, often-overlooked step by providing a data-driven method to justify the number of cut-points, protecting against arbitrary data dichotomisation. Furthermore, the package’s most unique contribution is the implementation of a genetic algorithm (method = “genetic”) to identify multiple cut-points simultaneously. As demonstrated with the rapeseed germination data, this feature is essential for uncovering complex, non-linear relationships between a biomarker and an outcome, such as the U-shaped effect of temperature on germination time, which standard dichotomisation methods would miss.

### Comparison with Existing Software

The development of *OptSurvCutR* builds upon a long history of tools designed for cut-point optimisation. Foundational graphical tools, such as *X-tile* (Camp et al., 2004), developed nearly 20 years ago, established the importance of log-rank-based optimisation by using an exhaustive systematic search to find up to two optimal cut-points. While pioneering, this approach was computationally limited to stratifying data into a maximum of three groups. Later, web applications like Cutoff Finder (Budczies et al., 2012), and GUI-based tools like *Evaluate Cutpoints* (Ogłuszka et al., 2019) made these analyses more accessible by wrapping existing R functions.

Within the current R ecosystem, *OptSurvCutR* complements and extends existing packages by providing a unique, integrated workflow (**Table 2**). While packages like *survminer* and *maxstat* are powerful tools for finding a single optimal cut-point, recent research has focused on more complex, non-linear scenarios. For instance, the *CutpointsOEHR* package implements a novel “optimal equal-HR” method specifically designed to find two cut-points in data with U-shaped relationships (Chen et al., 2019). The *Evaluate Cutpoints* application, in turn, uses a hierarchical method to find a second cut-point after an initial split (Ogłuszka et al., 2019).

**Table 2:** Functional and technical comparison of *OptSurvCutR* with other cut-point software. The table contrasts the key features of *OptSurvCutR* against other relevant R packages (*survminer, maxstat, CutpointsOEHR*), a GUI-based application (*Evaluate Cutpoints*), and the foundational bioinformatics tool (X-tile, Cutoff Finder). The compared features include: the primary Platform/User Interface; the core Multi-Cutpoint Method, where ‘Simultaneous’ optimisation is generally more robust than ‘Sequential’ approaches; the Max. No. of Groups the tool can create; the presence of a data-driven function for Cut-point No. Selection; an integrated function for Built-in Validation of cut-point stability, the algorithm’s flexibility for different relationship shapes (Generality of Application), and the ability to account for confounding variables (Covariate Adjustment).

| Feature | *OptSurvCutR* | *survminer* / *maxstat* | *CutpointsOEHR* | *Evaluate Cutpoints* | X-tile | Cutoff Finder |
| --- | --- | --- | --- | --- | --- | --- |
| **Platform / User Interface** | R Package / (GUI planned) | R Packages | R Package | Shiny App / R Package | Standalone GUI | Web App (Defunct) |
| **Multi-Cutpoint Method** | Simultaneous (Genetic Alg.) | N/A (1 cut only) | Constrained (Equal-HR) | Sequential (Hierarchical) | Systematic (Exhaustive) | N/A (1 cut only) |
| **Max. No. of Groups** | Unlimited | 2 | 3 | 3 | 3 | 2 |
| **Cut-point No. Selection** | Yes (Data-driven) | No | No | No | No | No |
| **Generality of Application** | High (any shape) | N/A (1 cut only) | Low (U-shape only) | Medium (wrapper-limited) | Medium | N/A (1 cut only) |
| **Built-in Validation** | Yes (Bootstrap) | No | No | No | No | No |
| **Covariate Adjustment** | Yes | No | Yes | No | No | No |

*OptSurvCutR* advances on these approaches by providing a more powerful and general-purpose solution. Its genetic algorithm finds all cut-points simultaneously, a method more likely to identify the global optimum compared to sequential approaches. Critically, it is not restricted to a specific relationship shape or a fixed number of thresholds. By combining this algorithmic flexibility with a complete workflow—including data-driven selection of the cut-point number find_cutpoint_number() and integrated bootstrap validation validate_cutpoint() — *OptSurvCutR* offers a more rigorous and comprehensive framework than is currently available in other specialised tools.

A key differentiator in the R ecosystem is the *method* used to find non-linear (e.g., U-shaped) cut points. As demonstrated in Case Study 2, *OptSurvCutR* successfully identifies these complex patterns by using its genetic algorithm in an unconstrained, statistics-first search. The algorithm’s only goal is to find the cut-points that best optimise the survival metric (e.g., the log-rank statistic), regardless of the underlying shape. This ‘agnostic’ approach is highly robust to ‘noisy’ real-world data and is not limited to U-shapes. This contrasts with specialised packages like *CutpointsOEHR*, which employ a constrained, model-first search that assumes a perfect U-shape *a priori* and analyses an idealised spline curve. While elegant, this ‘equal-HR’ method can be sensitive to data imperfections and is not a general-purpose discovery tool.

### The Importance of a Standardised Workflow

The development of *OptSurvCutR* also addresses the need for reproducible research in biomarker studies. By encapsulating these complex statistical procedures within a single, well-documented R package, we provide a transparent and standardised tool that enhances the reproducibility of cut-point analyses compared to the use of ad-hoc scripts. This emphasis on a robust, validated workflow aligns with guidance from health technology assessment, where understanding the assumptions and plausibility of a model’s long-term behaviour is critical (Rutherford et al., 2020). While ‘black box’ machine learning models may offer high predictive accuracy, their clinical utility is often hampered by a lack of interpretability. A primary advantage of *OptSurvCutR* is that it provides a complete workflow for generating interpretable and actionable risk models. The resulting outputs, such as Kaplan-Meier curves and validated thresholds, provide a clear ‘why’ behind a patient’s risk stratification, which is essential for clinical decision-making.

### Decoupling Model Selection from Cut-point Optimisation

A distinguishing feature of the OptSurvCutR workflow is the separation of model selection (determining the number of cut-points) from cut-point optimisation (determining their precise location). While it is possible to use the cut-points identified during the AIC/BIC search in the first step, we recommend a dedicated second optimisation step for two reasons.

First, statistical information criteria (AIC, BIC) prioritise model parsimony and likelihood maximisation, which may not always align perfectly with the clinical goal of maximising group separation (log-rank statistic) or effect size (hazard ratio). By establishing the optimal *number* of groups ($k$) using penalised criteria first, researchers avoid the “data dredging” pitfall of testing various $k$ values until a significant result is found.

Second, the genetic algorithms used (via rgenoud) are stochastic search methods. The search space for determining the number of cuts (e.g., 0 vs. 1 vs. 2) is distinct from the search space for optimising the location of a fixed number of cuts. A dedicated second step allows the algorithm to focus its computational resources entirely on refining the location of the $k$ cuts, often yielding slightly more precise boundaries for the chosen clinical metric.

### Bridging Machine learning Outputs and Clinical Utility

A valuable application for *OptSurvCutR* is in the post-hoc analysis and validation of prognostic scores generated by machine learning (ML) models. Such scores often require a clear threshold to be clinically actionable, yet common approaches like using the median are arbitrary and cohort-dependent, lacking a direct link to patient outcomes. Our workflow offers a principled approach to move beyond this by generating evidence for a potential clinical rule. First, the *find_cutpoint_number()* function allows researchers to objectively determine if the data truly supports stratifying patients into two or more groups, using information criteria like AIC or BIC to justify the model’s complexity. Next, *find_cutpoint()* identifies the precise location of this threshold by optimising a survival metric, such as the log-rank statistic, ensuring the cut-point is maximally associated with the outcome itself. Crucially, the workflow provides a framework to assess the stability of this finding using the bootstrap validation in *validate_cutpoint()*. A narrow 95% confidence interval from this analysis suggests the threshold is robust and less likely to be an artefact of the specific patient sample, whereas a wide interval serves as a warning of instability. This entire process can therefore serve as a crucial bridge, helping to translate an opaque ML score into a more interpretable and rigorously evaluated candidate rule for potential clinical use.

### Limitations and Future Directions

Despite its strengths, *OptSurvCutR* has limitations that present opportunities for future development. The primary algorithmic trade-off lies between the systematic and genetic search methods. The systematic search (method = “systematic”) guarantees a globally optimal solution by exhaustively evaluating all possibilities. Still, it suffers from a combinatorial increase in computational cost, making it impractical for finding more than two cut points in large datasets.

The genetic algorithm (method = “genetic”) provides an efficient heuristic alternative for these complex cases, but its stochastic nature means it is not guaranteed to find the global optimum in every run. It can get “converge” in a “local optimum,” returning a valid result that is statistically significant but suboptimal compared to the true best answer. Furthermore, its random starting population (pop.size) means it may, by chance, fail to find any valid solution on sparse data that does not meet the nmin constraint, even if a valid solution exists. The default settings are tuned for speed, but users can mitigate this limitation by increasing the search effort. Passing arguments for a larger max.generations (e.g., 500) to find_cutpoint forces a wider, more comprehensive search, making it much more likely to find the global optimum at the cost of computational time. We therefore strongly recommend method = “systematic” as the definitive method for 1-2 cut-points, reserving method = “genetic” for exploratory analysis or for 3+ cut-points where a systematic search is computationally infeasible.

Furthermore, from a statistical rigour perspective, while the bootstrap validation in *validate_cutpoint()* robustly assesses the stability of the chosen thresholds, the initial selection of the number of cuts could be strengthened. The addition of a formal permutation testing method to *find_cutpoint_number()* would provide a corrected p-value, offering a more rigorous assessment of the statistical significance of the chosen model complexity and further protecting against the multiple comparisons problem inherent in the search.

In terms of algorithmic expansion, incorporating other specialised methods from the literature would enhance the package’s versatility. For instance, implementing the “optimal equal-HR” method (Chen et al., 2019) would offer users a distinct, clinically principled approach for defining a “normal range” in U-shaped relationships, complementing the current methods that optimise for maximal statistical separation. Finally, to broaden the package’s impact beyond the R programming community, a key future direction is the development of a graphical user interface (GUI) using the R Shiny framework. This would provide an interactive, code-free platform, making the package’s robust workflow accessible to a wider audience of clinicians and biomedical researchers. Such an application could be bundled within the package for local use, ensuring data privacy and leveraging the user’s own computational resources, or deployed as a public web application.

### Conclusions

In conclusion, *OptSurvCutR* is a powerful and versatile R package that provides a complete, end-to-end workflow for optimal cut-point analysis in a time-to-event context. By integrating tools for determining the number of cuts, locating them using multiple structured algorithms, and validating the results, researchers can perform more rigorous, reproducible, and nuanced analyses across diverse scientific disciplines.

## DECLARATIONS

### Ethics approval and consent to participate

Not applicable.

### Consent for publication

Not applicable.

### Availability of data and materials

The *OptSurvCutR* package is freely available on GitHub at https://github.com/paytonyau/OptSurvCutR.

The rapeseed germination dataset used in Case Study 1 was computationally simulated. The R code to generate this dataset and to reproduce the complete analysis is provided within the package’s vignette, which is accessible via the command browseVignettes(package = “OptSurvCutR”).

## Supplementary information 1

### Generation of the germination Dataset

The germination dataset is a simulated dataset created to rigorously test the functionalities of the *OptSurvCutR* package, particularly its ability to identify multiple cut-points in data with non-linear relationships. The simulation process is grounded in the empirical findings reported by Haj Sghaier *et al*. (2022) on rapeseed germination and was conducted in two main parts.

#### Part 1: Simulation from Manuscript Parameters

The first part of the dataset was generated using parameters derived directly from the source manuscript. This included data for seven constant experimental temperatures (5, 10, 15, 20, 25, 30, and 35°C). For each temperature, we defined the key parameters that govern the germination process:

- **Time Window:** The *start_day* and *end_day* of the germination period.
- **Growth Rate:** The slope and intercept for a linear growth model.
- **Success Rate:** The final *germination_rate* observed at that temperature.

Using these parameters, a dataset was simulated to reflect the outcomes at these specific temperature points.

#### Part 2: Simulation from Interpolated Data

To provide a more continuous predictor variable and create a more challenging optimisation problem, a second dataset was simulated. This was achieved by linearly interpolating the key parameters (*slope, intercept, germination_rate*, etc.) for six intermediate temperature points (7.5, 12.5, 17.5, 22.5, 27.5, and 32.5°C) that were not present in the original study.

The final germination data object included in the package is the combination of these two simulated subsets. This hybrid approach yields a dataset that honours the original experimental findings while also providing the continuity needed to test and demonstrate cut-point finding algorithms robustly.

~~~
# ===================================================================
# DATA GENERATION SCRIPT for the ‘germination’ dataset.
# This script simulates rapeseed germination data based on parameters from
# Haj Sghaier et al. (2022) and saves the final object to data/germination.rda
# ===================================================================
# Set a global seed for full reproducibility of the simulation
set.seed(2025)
# ---1. Define Parameters & Core Simulation Function ---
# Original parameters from the manuscript
original_params <-data.frame(
  temperature = c(5, 10, 15, 20, 25, 30, 35),
  start_day = c(13, 7, 4, 4, 3, 4, 9),
  end_day = c(25, 20, 16, 16, 15, 16, 20),
  slope = c(0.9846, 3.6848, 4.9765, 3.8693, 6.5766, 1.1384, 0.1560),
  intercept = c(-13.627, -34.076, -28.116, -17.012, -35.621, -1.2833, -1.0571),
  r_squared = c(0.9585, 0.9501, 0.9307, 0.9524, 0.9564, 0.9237, 0.9574),
  germination_rate = c(0.88, 0.90, 0.92, 0.98, 0.94, 0.91, 0.55)
 )
# Core data generation function (unchanged)
generate_hybrid_data <-function(params) {
  set.seed(as.integer(params$temperature * 10))
  n_replicates <-4
  n_seeds_per_dish <-20
  n_samples_total <-n_replicates * n_seeds_per_dish
  time_data <-round(runif(n_samples_total, min = params$start_day, max = params$end_day))
  perfect_growth <-params$intercept + params$slope * time_data
  perfect_growth_variance <-var(perfect_growth)
  if (params$r_squared < 1 && params$r_squared > 0) {
  residual_variance <-perfect_growth_variance * (1 - params$r_squared) /params$r_squared
  } else
    { residual_variance <-0
  }
  residual_std_dev <-sqrt(residual_variance)
  residuals <-rnorm(n = n_samples_total, mean = 0, sd = residual_std_dev)
  actual_growth <-perfect_growth + residuals
  actual_growth[actual_growth < 0] <-0
  total_germinated <-round(n_samples_total * params$germination_rate)
  status_vector <-c(rep(1, total_germinated), rep(0, n_samples_total - total_germinated))
  shuffled_status <-sample(status_vector)
  replicate_vector <-rep(1:n_replicates, each = n_seeds_per_dish) actual_growth[shuffled_status == 0] <-0
  data.frame(
  temperature = params$temperature,
  replicate = replicate_vector,
  time = time_data,
  growth = actual_growth,
  germinated = shuffled_status
)
}
# ---2. Create Interpolated Parameters ---
# Define the new temperature points and the parameters to interpolate
new_temps <-c(7.5, 12.5, 17.5, 22.5, 27.5, 32.5)
params_to_interp <-c(“slope”, “intercept”, “r_squared”, “germination_rate”, “start_day”, “end_day”)
# Use sapply to apply the interpolation to all specified parameters at once
interpolated_values <-sapply(params_to_interp, function(param) {
approx(original_params$temperature, original_params[[param]], xout = new_temps)$y
})
interpolated_params <-as.data.frame(interpolated_values)
interpolated_params$temperature <-new_temps
# ---3. Combine Parameters and Run Full Simulation ---
# Combine original and interpolated parameters into a single data frame
all_params <-rbind(original_params, interpolated_params)
# Apply the generation function to every row of the combined parameter set
all_data_list <-lapply(1:nrow(all_params), function(i) generate_hybrid_data(all_params[i, ]))
# Combine the list of data frames into one final object named ‘germination’ germination <-do.call(rbind, all_data_list)
# ---4. Save the Final Data Object for the Package ---usethis::use_data(germination, overwrite = TRUE)
~~~

